# Identification roles of NFE2L3 in Digestive System Cancers

**DOI:** 10.1101/2023.12.05.570235

**Authors:** Fan Li, Zhili Wen

## Abstract

**(1) Background:** The morbidity and mortality rate of Digestive System Cancers (DSC) continue to threaten human lives and health. Nuclear factor erythroid 2-like protein 3 (NFE2L3) is associated with the development, growth, and progression of multiple cancers. Nevertheless, the clinical value and underlying mechanisms of NFE2L3 in DSC remains unknown. This study aimed to clarify the possibilities of NFE2L3 as a novel biomarker in DSC.

**(2) Methods:** We obtained the data of NFE2L3 in cancers from database to evaluate the expression level and clinical value of NFE2L3 in DSC, and to explore underlying mechanism and biological functions of NFE2L3 in DSC.

**(3) Results:** NFE2L3 expression is up-regulated in DSC and have both prognostic and diagnostic value. NFE2L3 contributes to oncogenesis through a variety of mechanisms. In vitro experiment showed that NFE2L3 stimulated the proliferation and migration ability of gastric cancer cells.

**(4) Conclusion:** Our study confirms the clinical applicability of NFE2L3 as a promising biomarker for DSC.

## 1. Introduction

Globally, Tumors continue to pose a threat to human life and health. In 2021, statistics from the American Cancer Society showed that there were 338090 new cases and 169280 deaths were estimated for digestive tract tumors in the United States ^[1]^. DSC mainly include cholangiocarcinoma (CHOL), colon adenocarcinoma (COAD), esophageal carcinoma (ESCA), liver hepatocellular carcinoma (LIHC), pancreatic adenocarcinoma (PAAD), rectum adenocarcinoma (READ), stomach adenocarcinoma (STAD), continues to be a major type of cancer threatening human health and survival. DSC are affected by several risk factors, including lifestyle habits, tobacco use, alcoholism, physical activity, and gender ^[2]^. Therefore, it is of great importance to identify a biological target with ideal diagnostic and therapeutic importance for patients with tumors.

NFE2L3 was discovered in the 1990s, belonging to the cap ’n’ collar basic-region leucine zipper family, mainly located in the endoplasmic reticulum and the nuclear envelope ^[3]^. Currently, researches related toap ’n’ collar basic-region leucine zipper family primarily focuses on nuclear factor erythroid-derived 2-like 2 (NFE2L2). NFE2L2 plays the role of the primary transcriptional regulator for the cell’s antioxidant response controls, which are associated with cancer and drug resistance. The binding of kelch like ECH associated protein 1 (KEAP1) to NFE2L2 causes polyubiquitylation, leading to swift degradation of NFE2L2 and significantly influencing the initiation, progression, and evolution of tumors. The mechanism of NFE2L3 in cancers is entirely different from NRF2, but limited research has evaluated the significance of NFE2L3 as an innovative indicator in malignancies.

NFE2L3, as a crucial regulator of the cellular stress response, expressed in diverse tissues of the human, different levels of expression highlight the unique functionality of NFE2L3 in different tissues ^[4–6]^. In recent years, a growing number of evidence has indicated the participation of NFE2L3 in the processes of differentiation, inflammation, and carcinogenesis. James Saliba demonstrated that the deletion of NFE2L3 results in decreased inflammation and a shift in the tumor microenvironment via the IL33 and RAB pathways ^[7]^. Knockdown of NFE2L3 inhibited bladder cancer cell proliferation by inducing the cell cycle and apoptosis ^[4]^. In vitro and in vivo experiments support NFE2L3 as an oncogene involved in the process of tumor formation. Although findings have suggested that NFE2L3 may be a crucial regulator of cancer progression, the underlying molecular mechanisms remain poorly understood.

The prevalent approach to treating cancer involves a combination of chemotherapy, radiotherapy, and surgical excision of the tumor. Tumor immunotherapy as a novel therapeutic modality has revolutionized cancer treatment ^[8–10]^. The investigators discovered that NFE2 family is a potential prognostic biomarker and is associated with immune infiltration in many cancers ^[11]^.

The goal of the study was to investigate the possibility of NFE2L3 as novel biomarker for DSC. We import data from different database to provide a systematic and comprehensive insight into NFEE2L3 expression level, diagnostic and prognostic value, and underling mechanism in DSC, we also used patients specimens and *in vivo* models test to confirm the biological functions of NFE2L3 in DSC.

## 2. MATERIALS AND METHODS

### 2.1 Gene Expression Analysis

All raw data were downloaded from The Cancer Genome Atlas (TCGA), University of California Santa Cruz (UCSC) Xena browser (https://xena.ucsc.edu/), and the Human Protein Atlas (HPA) (https://www.proteinatlas.org/). Raw data were processed using Strawberry Perl 5.30.0.1 and R 4.1.1. The Wilcoxon test was used to analyze the data. p < 0.05 (*) was considered statistically significant.

Twelve gastric cancer and normal tissues from patients admitted to Second Affiliated Hospital of Nanchang University. Written informed consent was obtained from the patients. This study was conducted in accordance with the guidelines of the Declaration of Helsinki, and approved by the Medical Ethics Committee of the Second Affiliated Hospital of Nanchang University (No. 2022096 Approve data: 2022.9.7).

### 2.2 Tumor mutational burden and microsatellite instability of NFE2L3 in pan-cancer

We extracted relevant data of Tumor mutation burden (TMB) and Microsatellite Instability (MSI) fro raw data, and the relationship between NFE2L3 and pan-cancer were analyzed using Spearman’s rank correlation coefficient.

### 2.3. Clinical relevance of NEF2L3 in DSC

We created receiver operating characteristic (ROC) curves using the R package "pROC" to examine the diagnostic value of NEF2L3 in DSC. The ROC curve area (AUC) between 0.7 and 0.9 indicated sure precision, and the AUC over 0.9 indicated great accuracy.The prognostic effect of NFE2L3 in DSC was investigated by performing Kaplan-Meier survival analysis. We selected three markers for the Kaplan-Meier analysis: overall survival (OS), disease-specific survival (DSS), progression-free interval (PFI). Both the "survival" and "survminer" packages were used to plot the survival curves. The relationship between NFE2L3 expression and clinical features, using "limma" and "ggpubr" packages.

### 2.4 NFE2L3 expression in subtypes and immune imfiltration

NFE2L3 expression in immunological or molecular subtypes were measured in TISIDB (http://cis.hku.hk/TISIDB/), which was a website for tumor immune interaction. The immune cell composition of DSC was analyzed using CIBERSORT (https://cibersort.stanford.edu/). R packages “ggplot2”, “ggpubr” and “ggExtra” were used to process the data and calculated the correlation coefficient. |R|≥0.3 and *p* <0.05 (*) were considered statistically significant.

### 2.5 Co-expression genes of NFE2L3 in DSC

We using the packages "limma" and "BiocManager" to investigate the co-expressed genes and chemokine receptors that were both positively and negatively related to NFE2L3 expression in DSC.

### 2.6 Protein-protein interaction Network building

A protein-protein interaction (PPI) network was constructed using Search Tool for the Retrieval of Interacting Genes (STRING) (https://string-db.org/), with the following fixed principal parameters: a minimum required interaction score of 0.4, a maximum of 50 inter actors to be displayed, and the sources of active interaction from experiments, text mining, and databases. The PPI networkwas visualized using Cytoscape (version 3.9.0).

### 2.7 Gene Ontology and Kyoto Encyclopedia of Genes and Genomes pathway enrichment analysis

Gene Ontology (GO) and Kyoto Encyclopedia of Genes and Genomes (KEGG) enrichment analyses and Genomes Enrichment Analyses (GSEA) were performed to assess the potential biological functions and pathways of NFE2L3 in DSC, using the packages “clusterProfiler”, “ggplot2” and “limma”.

### 2.8 qRT-PCR

Total RNA extraction was extracted using the TRIzol Up Plus RNA Kit (Trnsgen). The cDNA synthesis of cDNA using M-MLV reverse transcription (Trnsgen). Quantitative reverse transcription polymerase chain reaction (qRT-PCR) was performed by SYBR Green PCR Master Mix (Takara). The primer sequences for qRT-PCR were listed in Table S15.

### 2.9 Cell line and culture

HGC-27 and AGS were purchased from the American Type Culture Collection (ATCC). Cells were cultured in RPMI-1640 containing 10% fetal bovine serum and cultured at 37℃ containing 5% CO2.

### 2.10 siRNAs and Plasmids Transfection

Plasmids (pcDNA3.1)were purchased from Shanghai Genechem. Plasmids were transfected into AGS via Lipofectamine 2000 (ThermoFisher). siRNA (siNFE2L3#1, siNFE2L3#2, siNFE2L3#3) purchased from RIBOBIO and transfected into HGC-27 using Lipofectamine 2000. qRT-PCR was applied to detect the efficiency of NFE2L3 overexpression or knockdown. The sequences of siRNA were listed in Table S16.

### 2.11 Proliferation and migration assays

For proliferation experiments, approximately 1000 cells per well for each cell line were seeded in 96-well plates and Tthe 450nm absorbance was tested for 6 days (0, 1, 2, 3, 4, 5 days). The CCK-8 reagent was purchased from APExBIO (USA). We seeded 3×10^5^ cells per well in 6-well plate and collect images at 0h, 12h and 24 h to calculate migration rates over time.

### 2.12 Statistical Analysis

The statistical analysis in this study was automatically calculated using the aforementioned online databases. *p* value <0.05 was checked as the statistical significance. \**p*<0.05, \*\**p*<0.01, \*\*\**p*<0.001.

### 3. Results

### 3.1 NFE2L3 Expression Level in Pan-Cancer

To identify the expression level of NFE2L3 in cancers, a comparison is made to the normal tissue and cancer tissues, the results showed that NFE2L3 expression levels were elevated in DSC (Figure 1A, B). Results were further validated by quantitative real-time PCR (qRT-PCR) in twelve STAD samples and its cancer adjacent tissues (Figure 1F). The clinical characteristics of patients were presented in the supplemental materials (Table S14). The data from the Human Protein Atlas (HPA) annotate the abnormal expression of NFE2L3 in most tumor cell lines and cancer tissues (Figure 1C, D, E).

**Figure 1.**
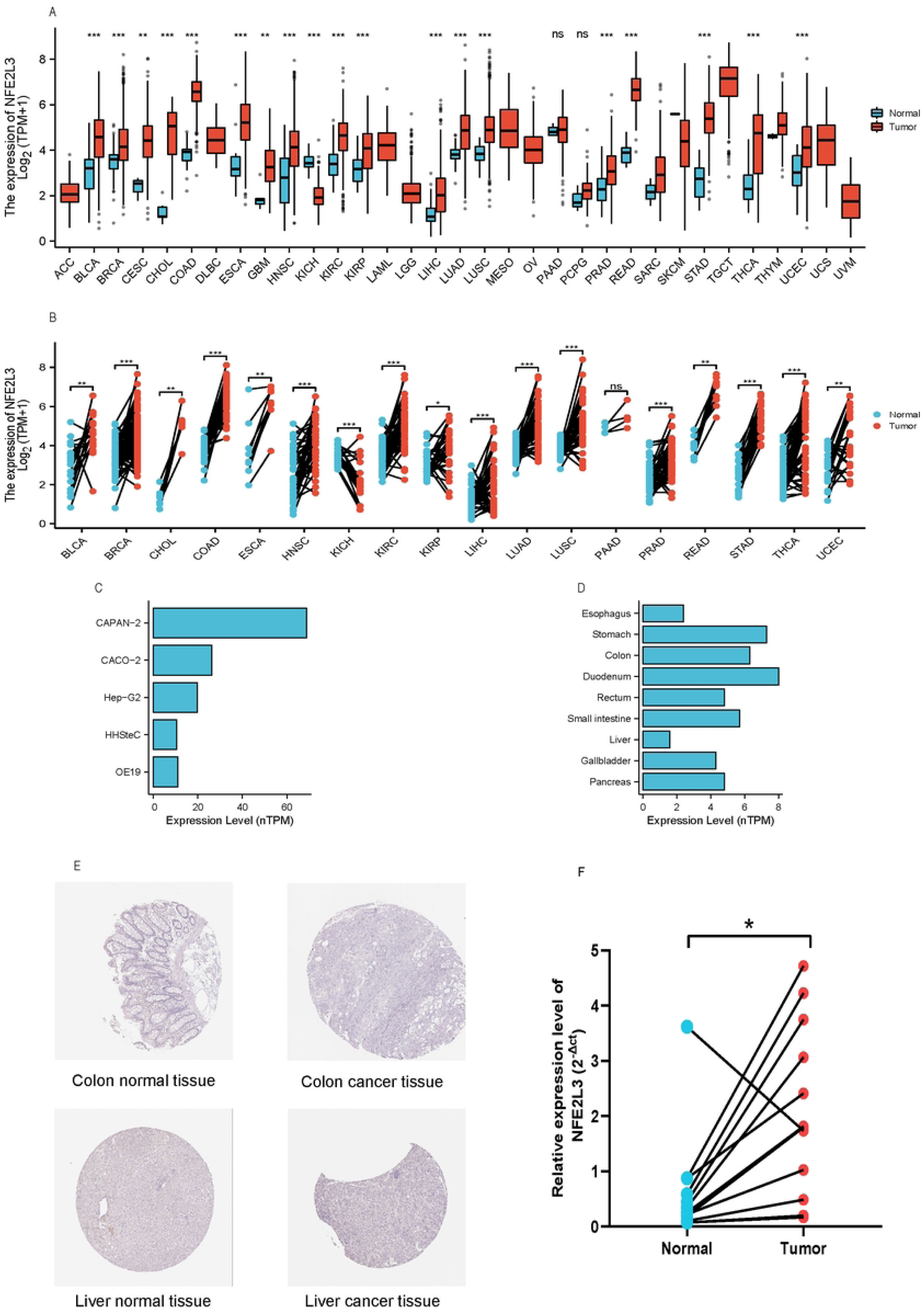
Expression level of NFE2L3 A. NFE2L3 expression in pan-cancer; B. NFE2L3 expression in tumor tissues and paracancerous tissues; C. NFE2L3 expression in DSC cell lines; D. NFE2L3 expression in DSC tissues; E. Immunohistochemical staining of colon and liver tissues; F. NFE2L3 expression in gastric cancer and normal tissues.

### 3.2 TMB and MSI of NFE2L3 in DSC

MSI were observed in STAD, THYM, COAD, DLBC, SKCM, and LUAD (Figure 2A). TMB occured in ACC, BLCA, BRCA, LAML, LGG, PAAD, and STAD, COAD, KIRC and UCEC (Figure 2B). The data implies that the MSI burden and TMB levels are important hallmark of tumor response of DSC to immune checkpoint blockade. The *p* values and the relevant coefficients of MSI and TMB are shown in the supplementary materials (Table S1, S2) .

**Figure 2.**
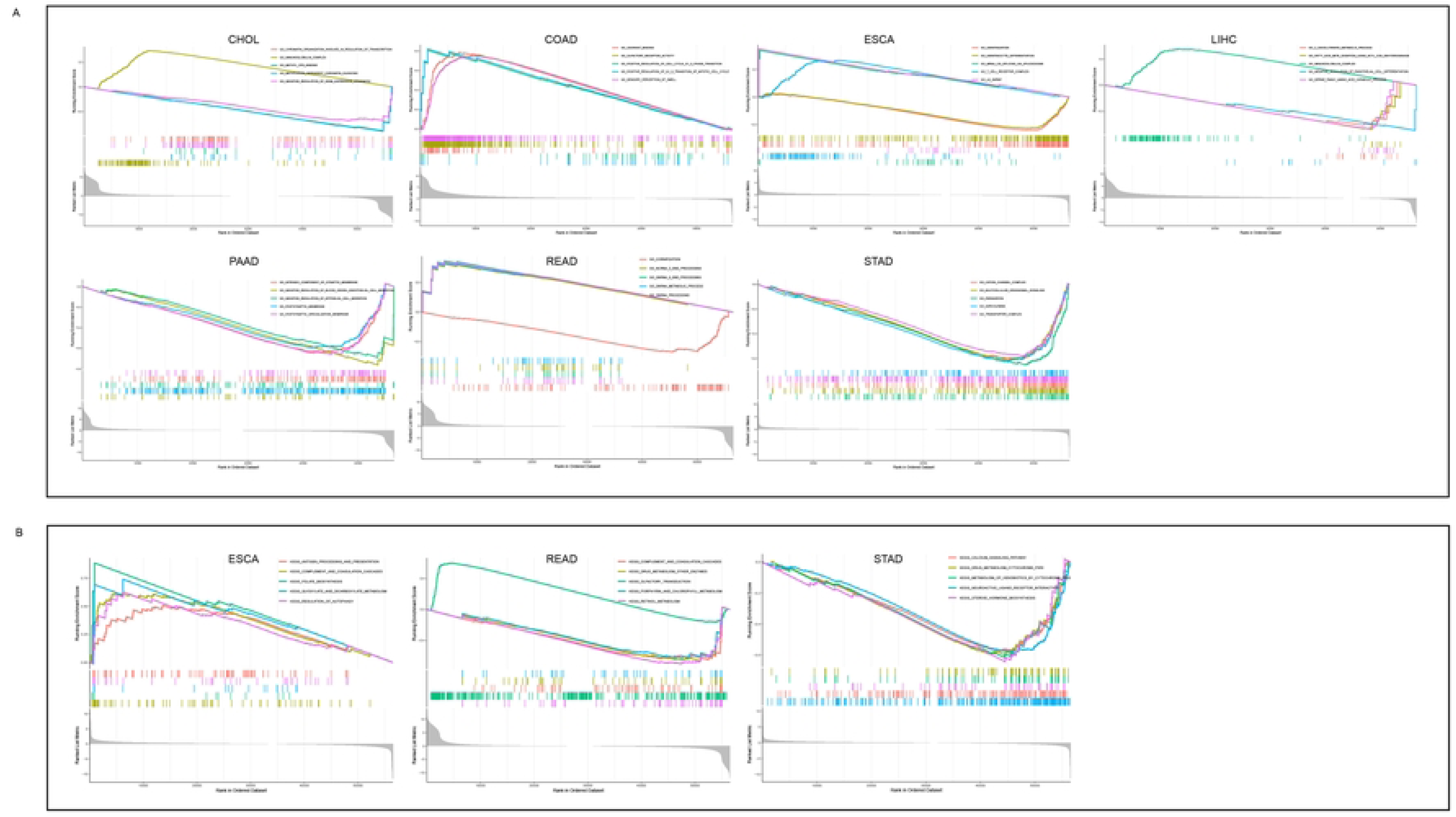
Relationship between NFE2L3 and MSI or TMB in DSC A.MSI; B.TMB

### 3.3. Diagnostic value of NFE2L3 in DSC

Based on the ROC curves, we concluded that NFE2L3 has a potential diagnostic value for DSC (Figure 3), particularly in STAD. The AUC values of ESCA, PAAD, LIHC, READ, and STAD were in the range of 0.87 to 1.00, indicating an ideal diagnostic value of NFE2L3 in theabove-mentioned cancers.

**Figure 3.**
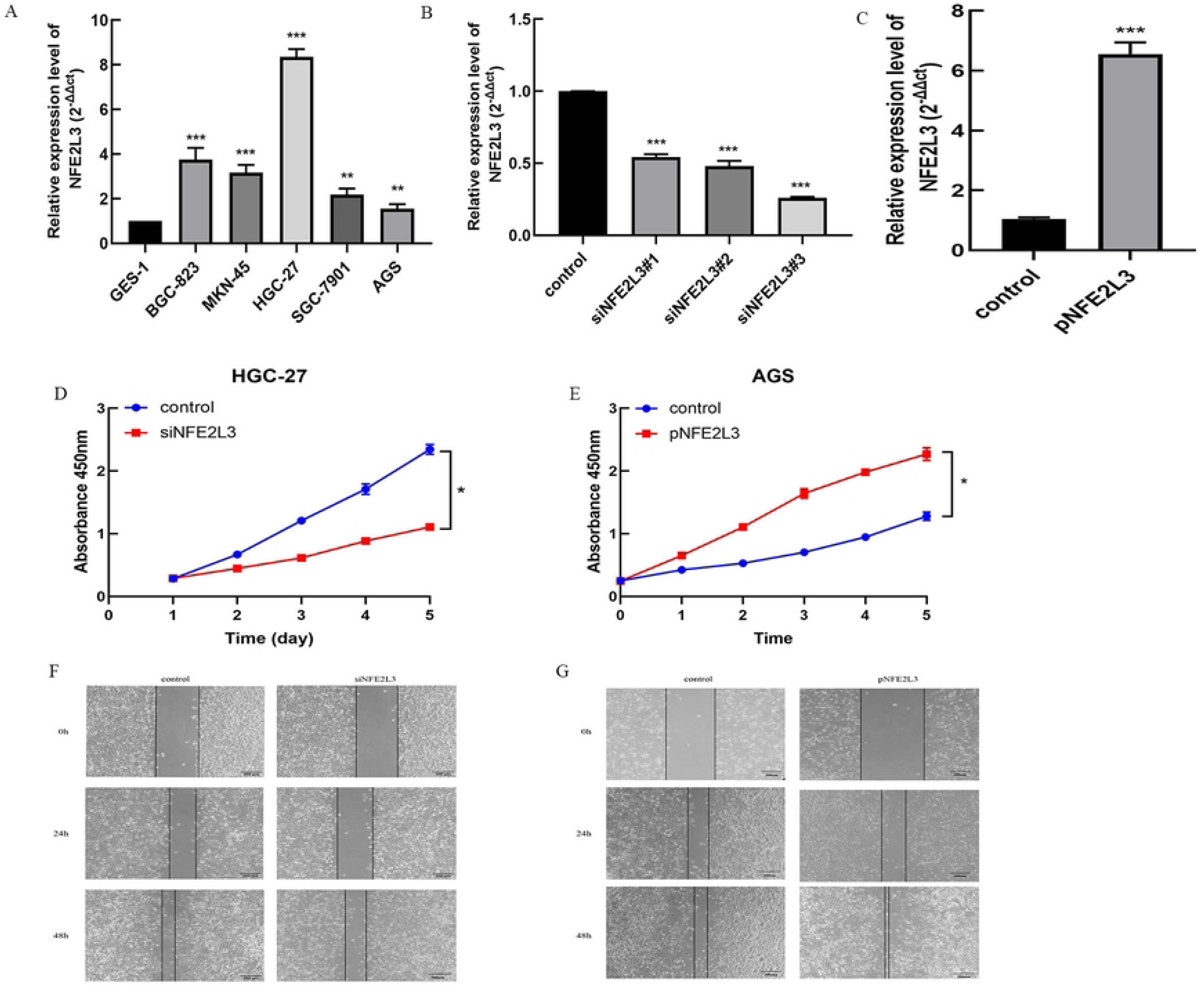
ROC curve of NFE2L3 in DSC A. CHOL; B. COAD; C. ESCA; D. PAAD; E. LIHC; F. READ; G. STAD

### 3.4. Prognostic value of NFE2L3 in DSC

The prognostic value of NFE2L3 in DSC were analyzed by comparing the overall survival (OS), progress free interval (PFI) and disease specific survival (DSS). In PAAD, high expression of NFE2L3 associated with poor OS, PFI and DSS. Lower NFE2L3 expression was associated with higher PFI in LIHC. These findings indicate that NFE2L3 could be identified as a prognostic indicator.

**Figure 4.**
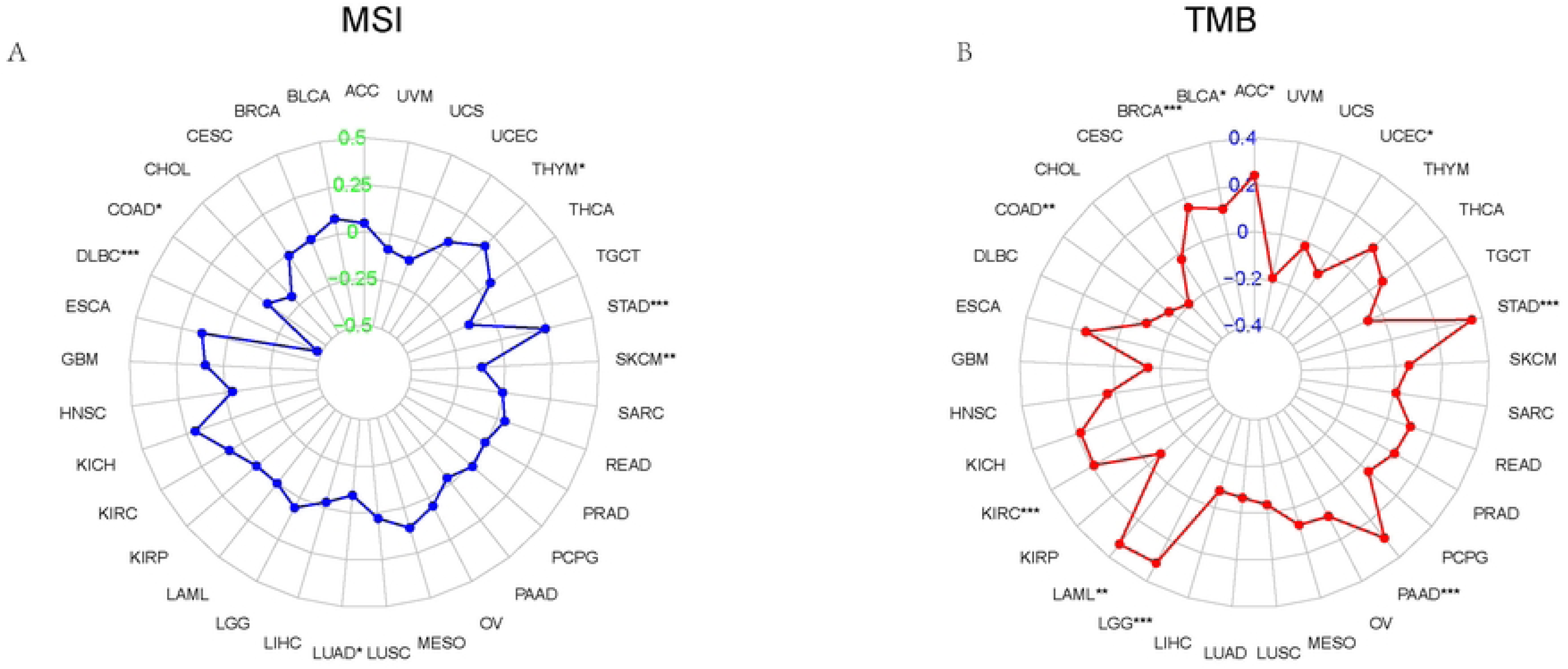
Correlations between NFE2L3 expression and prognosis A. OS of PAAD; B.PFI of PAAD; C. DSS of PAAD; D. PFI of LIHC

### 3.5. Clinical characteristics of NFE2L3 in DSC

To examine potential associations between clinical features and NFE2L3, we performed correlation analysis. There was a substantial correlation between NFE2L3 expression and both the T (*p*=0.006) and N (*p*=0.032) stages in COAD. Age was substantially correlated with NFE2L3 expression in patients with ESCA (*p*=0.012). In LIHC, NFE2L3 was strongly correlated with age (*p*=0.034), T stage (*p*=0.002) and pathological stage (*p*=0.001). The pathogenic stage of PAAD was substantially correlated with NFE2L3 expression level (*p*=0.017). There was a substantial correlation between the pathological stage of STAD and NFE2L3 expression (*p*=0.032). The clinical features of NFE2L3 in DSC are displayed in the supplemental materials (Table S3-S9). Results presented above suggest that NFE2L3 has a close association with clinical characteristics of patients.

### 3.6. NFE2L3 Expression in immune and molecular subtype

We observed both tumor immune (Figure 5) and molecular subtypes (Figure 6) of NFE2L3 in DSC. In READ, LIHC and CHOL, NFE2L3 was predominantly expressed in the C1. In STAD and PAAD, NFE2L3 was expressed in the C2. NFE2L3 expressed in C4 in ESCA and COAD. In terms of the molecular subtype, NFE2L3 was expressed in the CIN of the COAD and ESCA. LIHC showed that NFE2L3 was mainly expressed in iCluster1. In READ, NFE2L3 was primarily expressed in the GS molecular subtype. In STAD, NFE2L3 was primarily expressed in the molecular subtype of the HM-indel. The results showed that NFE2L3 presents with multiple molecular and immune subtypes in DSC, thereby making NFE2L3 suitable targets for immunotherapy.

**Figure 5.**
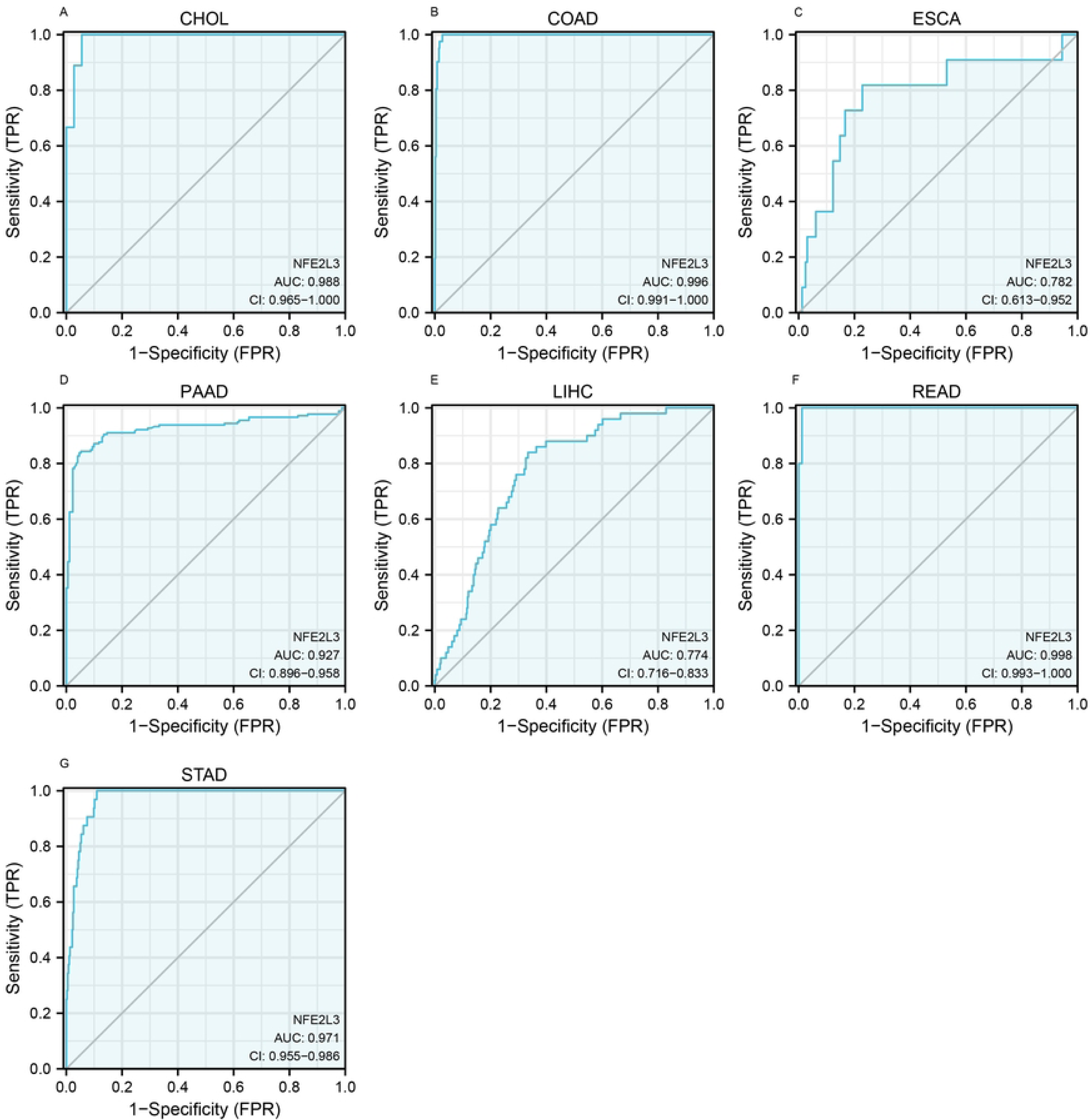
Correlations between NFE2L3 expression and immune subtypes A. PAAD; B. READ; C. LIHC; D. STAD; E. ESCA; F. CHOL; G. COAD

**Figure 6.**
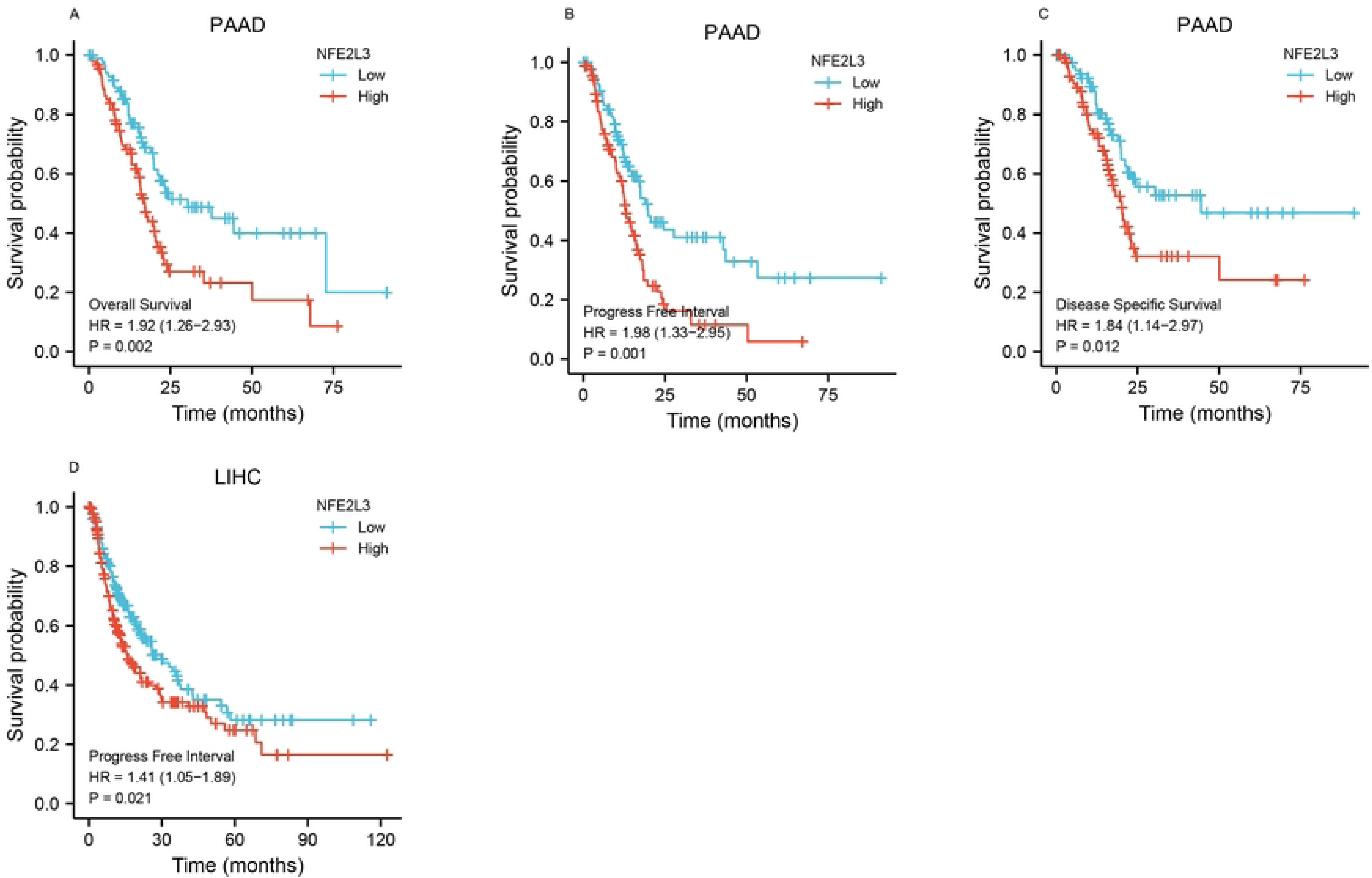
NFE2L3 expression and tumor immune in DSC A. COAD; B. ESCA; C. READ; D. LIHC; E. STAD

### 3.7 NFE2L3 expression and immune-related cell infiltration

We examined the relationship between NFE2L3 expression and immune-related cell infiltration in DSC (Figure 7). In LIHC and STAD, we found strong correlations between NFE2L3 expression levels and different immune cells, especially in T-regulatory cells (Tregs) and CD4^+^ T cells. This result suggested that there was close correlation between immune infiltration and NFE2L3 expression, NFE2L3 has the potential to be an important target for immunotherapy for DSC.

**Figure 7.**
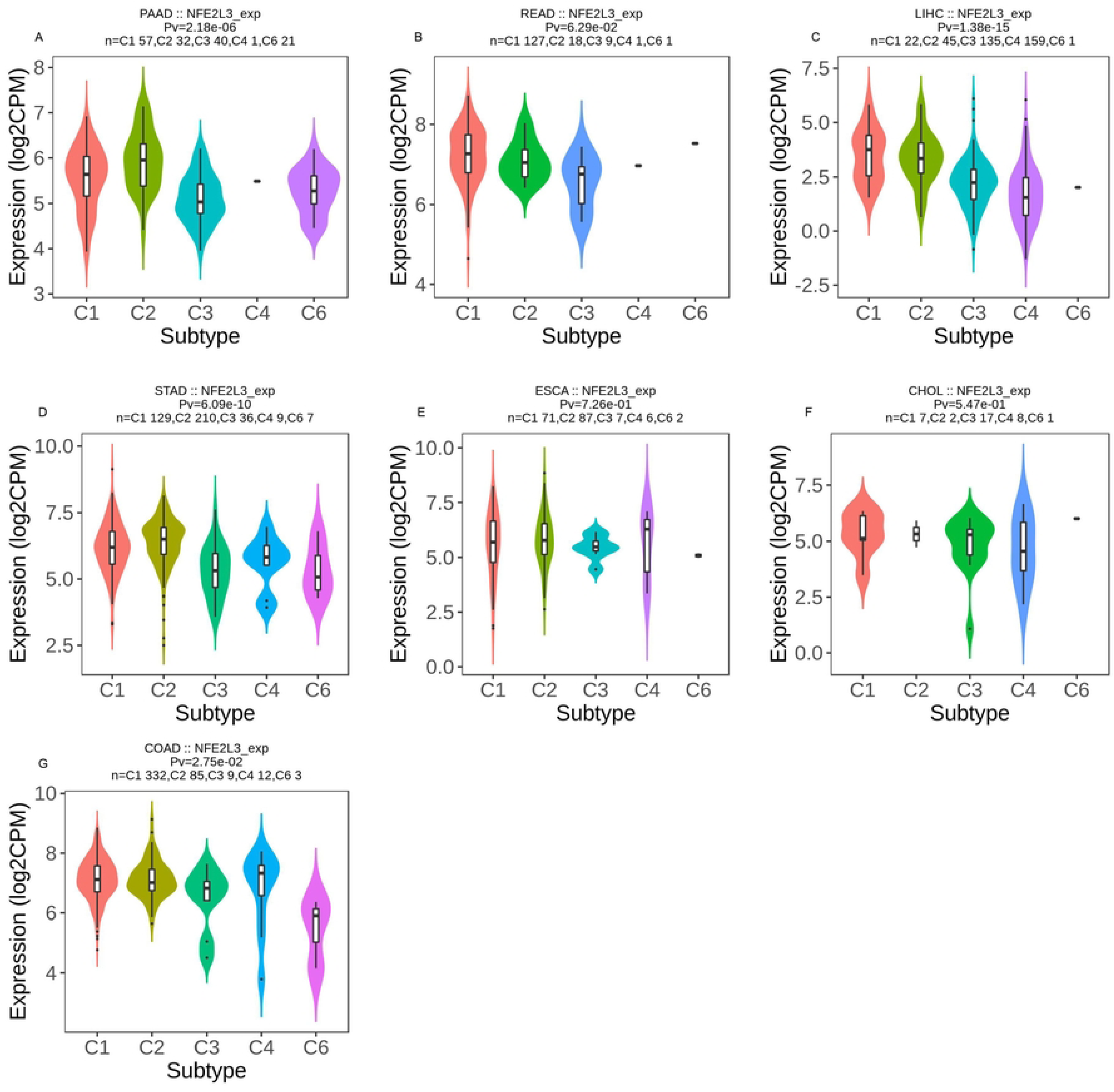
NFE2L3 expression and tumor immune in DSC A. COAD; B. ESCA; C. READ; D. LIHC; E. STAD

3.8 NFE2L3 in immune-related genes and chemokine receptors

In this study, we aimed to determine the roles of NFE2L3 in immune related genes (Figure 8A) and chemokine receptors (Figure 8B). The results showed that NFE2L3 presenting a positive correlation with the majority of immune related genes in LIHC, such as CD200 (*p*=0.00, R=0.339), CD274 (*p*=0.00, R=0.396), and CTLA4(*p*=0.00, R=0.475). Many genes, such as TNFRSF9, CD274, CD44, TIGIT, TNFSF15, TNFRSF18, TNFRSF4, VISR, TNFSF9, IDO1, VTCN1, CD80, TNFRSF14 and CTLA4 exhibit significantly abnormal expression in most DSC. The correlation analysis of co-expression between NFE2L3 and chemokine receptors showed that numerous chemokine receptors, such as CX3CR1, CXCR4, ACKR1, PTGDR2, LTB4R, and PLXNB2 shared close co-expression relationships with NFE2L3 in most DSC, most notably in LIHC, COAD and STAD. The related coefficients and *p* value are reported in the supplemental materials (Table S10-13).

**Figure 8.**
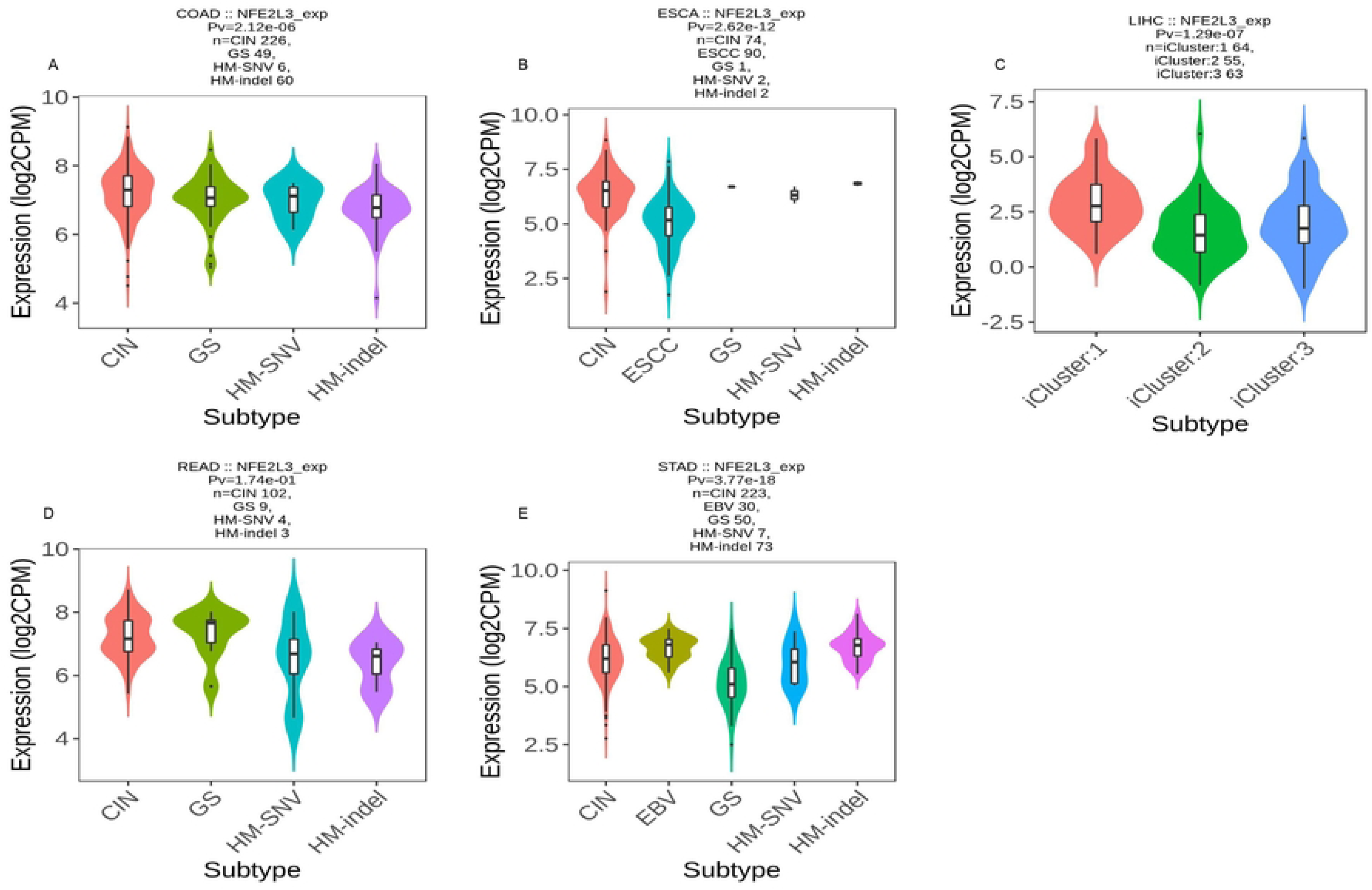
The relationships between NFE2L3 and immune-related genes or chemokine receptors A. Immune-related genes; B. Chemokine receptors

### 3.9 Proteins interact with NFE2L3

To obtain the potential target genes of NFE2L3, we conducted a PPI network. The results revealed that MAFK, NFE2, KEAP1, NFW2L2, MAFG, CUL3, HMOX1, NQO1, MAF, and GCLC are strong candidate target genes of NFE2L3. (Figure 9A). To further interrogate the mechanism of NFE2L3 interaction, we examined its interaction through Go and KEGG analysis. The GO analysis results (Figure 9B) showed that the activity of DNA-binding transcription, specificity of RNA polymerase II, and catabolic processes of proteaosomal proteins were the major functions of the interactions. The KEGG analysis revealed that the protein pathways were mostly focused on the hedgehog signaling pathway (Figure 9C).

**Figure 9.**
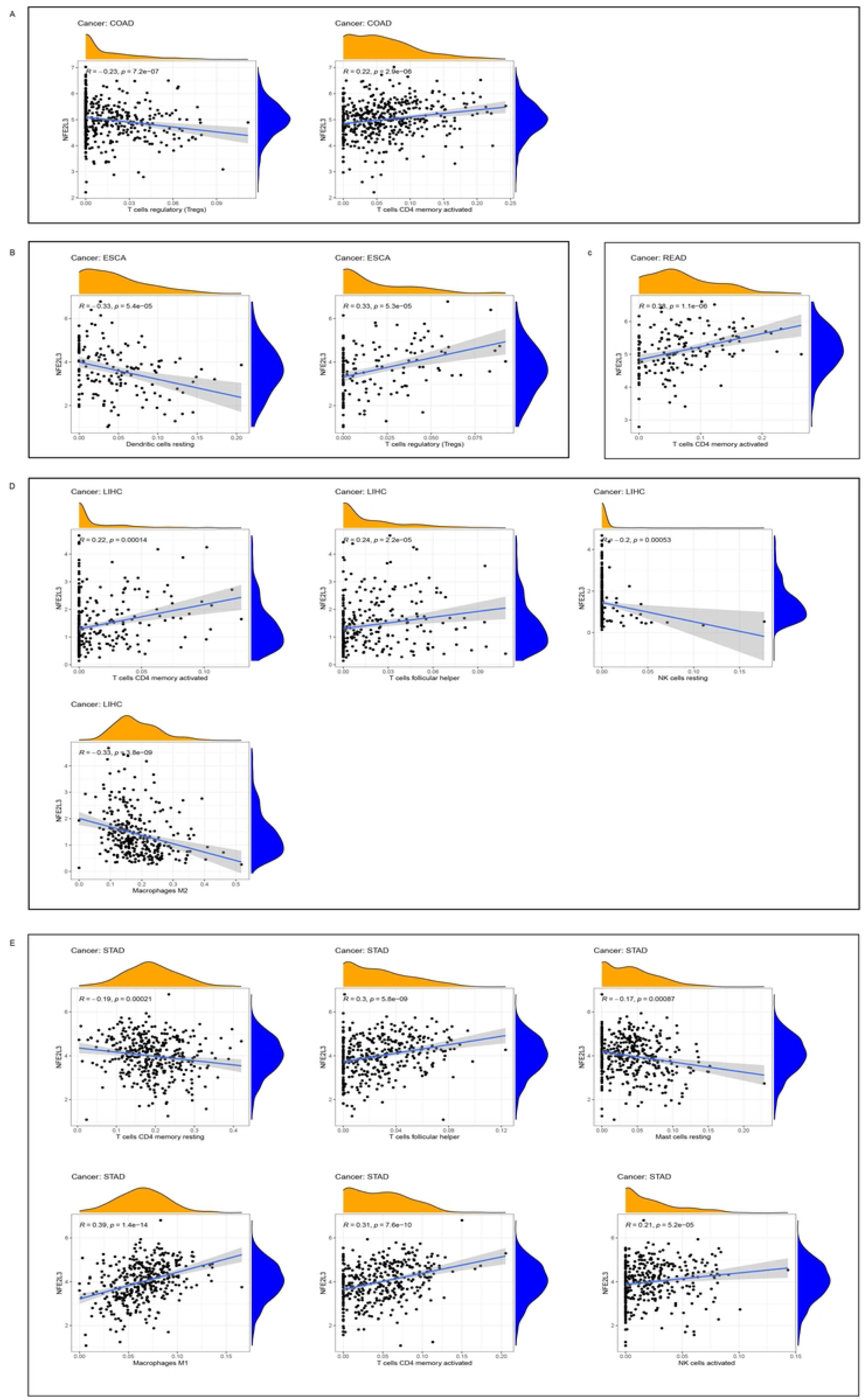
Proteins interact with NFE2L3, Go analysis and KEGG analysis of binding protein A.Proteins interact with NFE2L3; B. Go analysis of binding proteins; C. KEGG analysis of binding proteins

### 3.10 Gene Set Enrichment Analysis

To examine the biological significance of NFE2L3 expression in DSC, we conducted GSEA. The TCGA tumor samples were categorized into two groups: high- and low-expression groups. The top GO terms and KEGG pathways were analyzed using GSEA. The GO annotation results (Figure 10A) indicated that NFE2L3 negatively regulated methylation in COAD and the metabolic processes in LIHC. In STAD, NFE2L3 expression affected signal transduction. The KEGG terms (Figure 10B) indicated that NFE2L3 activated autophagy in ESCA, whereas it inhibited drug metabolisms, such as P450, as well as calcium signaling pathways in READ and STAD.

**Figure 10.**
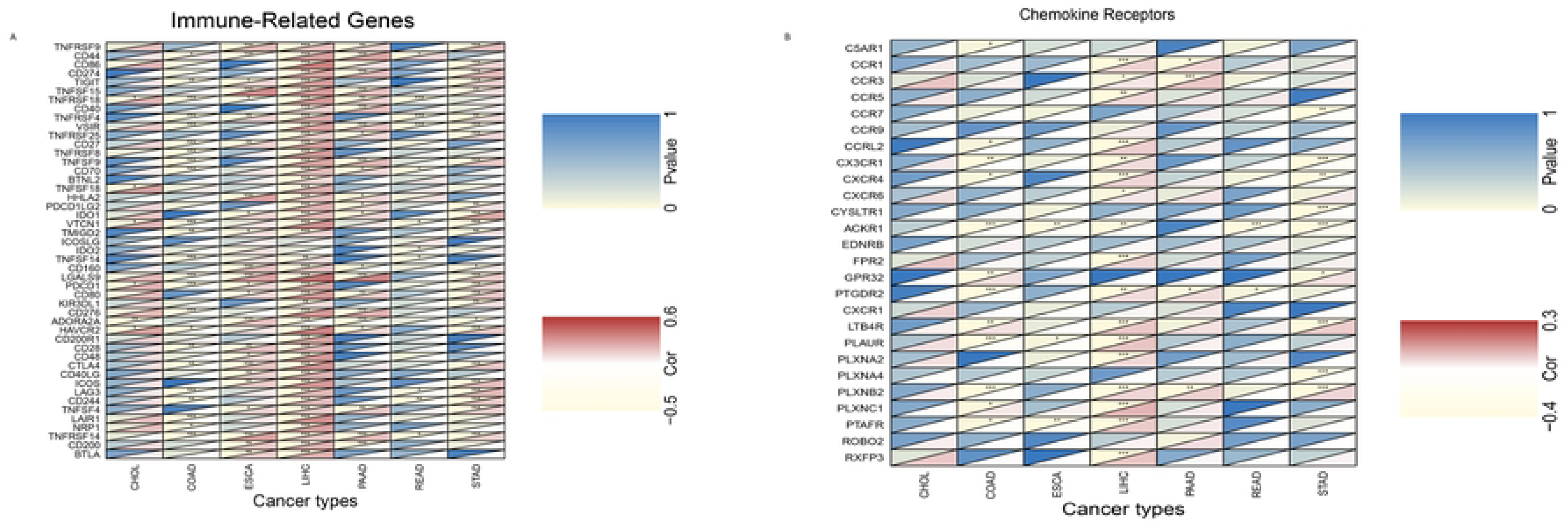
GSEA of NFE2L3 expression in DSC A.Go analysis; B. KEGG analysis

### 3.11 Biological functions of NFE2L3

NFE2L3 expression level in gastric cancer cells was observed to be higher compared with normal gastric mucosal epithelial cells (Figure 11A). Plasmids and siRNAs have an ideal efficacy in altering NFE2L3 expression levels (Figure 11B-C). CCK-8 and wound healing assay showed that NFE2L3 knockdown suppressed the proliferation and migration of HGC-27. At the same time, overexpression of NFE2L3 promotes the cellular proliferation, migration and invasion of AGS (Figures 11D-G).

**Figure 11.**
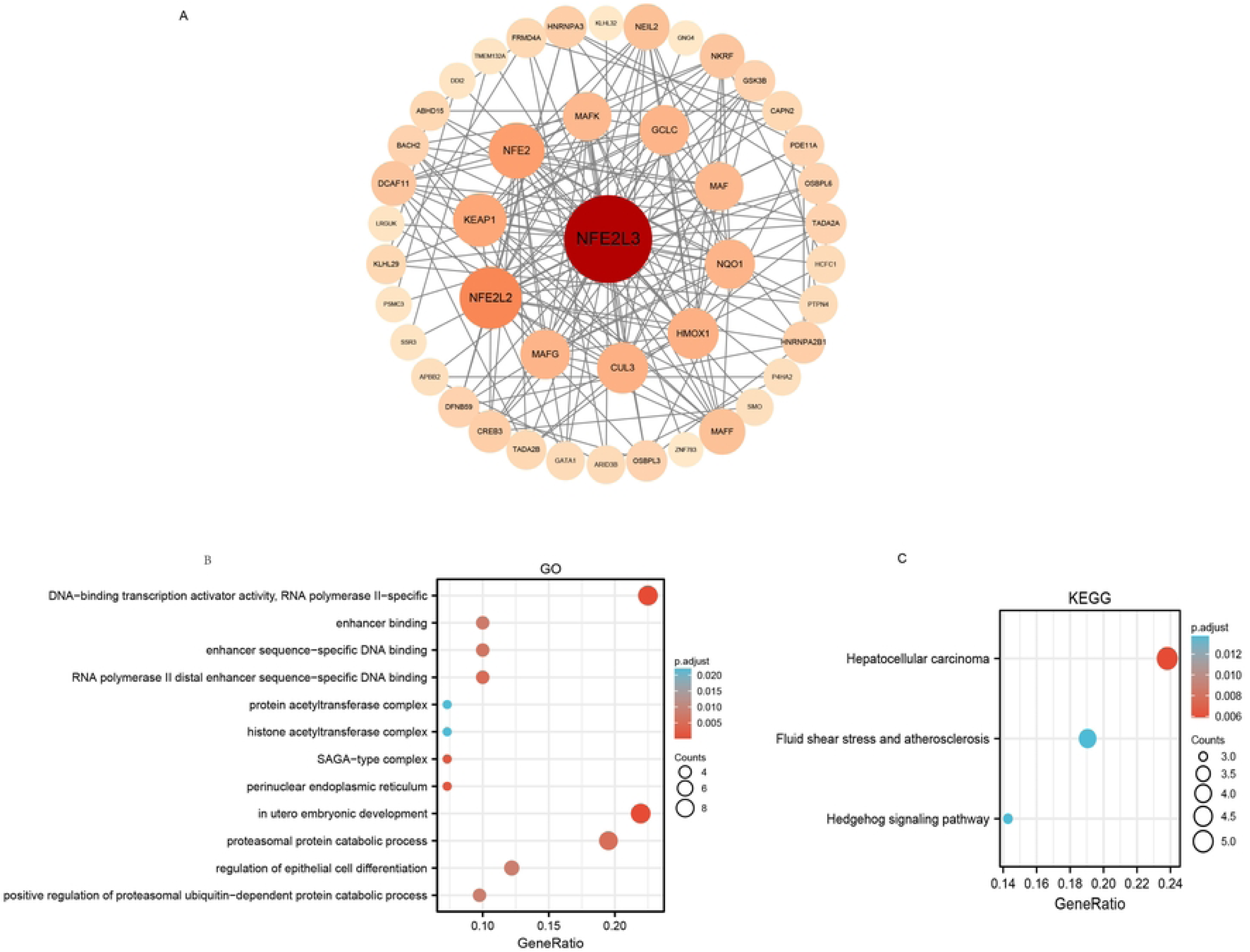
The biological function of NFE2L3 in gastric cancer A. The expression levels of NFE2L3 in cell lines; B. Verification of knockdown efficiency of NFE2L3 in HGC-27; C. Verification of overexpression efficiency of NFE2L3 in AGS; D. The proliferation ability of NFE2L3 in HGC-27; E. The proliferation ability of NFE2L3 in AGS; F. The migration ability of NFE2L3 in HGC-27; G. The migration ability of NFE2L3 in AGS.

## 4. Discussion

Our work in this study provides a comprehensive insight into the importance of NFE2L3 in DSC and revealed that NFE2L3 has great potential to become a widely utilized biomarker. NFE2L3 is found to be abnormally expressed in cancers and regulates DSC through multiple mechanisms. NFE2L3 has ideal clinical relevance for its good prognostic and diagnostic value.

Accumulating evidence suggests that NFE2L3 is crucial for the development of cancer. NFE2L3 belongs to the Cap‘n’ Collar family, which plays a vital role in the cancer-promoting function by various mechanisms ^[12, 13]^. For exampe, NFE2L3 functions as an oncogene by triggering the Wnt/-catenin signaling pathway ^[14]^, absence of the NFE2L3 affects the formation and growth of tumors ^[10, 11, 15]^. NFE2L3 regulates the immune microenvironment and has potential prognostic value in patients with kidney cancer ^[10]^. The absence of NFE2L3 results in significantly less inflammation in the colon, which inhibits an increase in the number and volume of tumors ^[11]^. Although the close relationship between NFE2L3 and cancers has been demonstrated, the clinical value and underling menchanism of NFE2L3 in cancers remains unclear.

A pan-cancer analysis showed that NFE2L3 is abnormally expressed in most malignancies, predominantly in the DSC. These findings have also been confirmed in cell lines and in cancer tissue data obtained from the HPA database. We further confirm this speculation in gastric cancer tissues and cell lines, the qRT-PCR results showed significant levels of NFE2L3. The above results lead to a important conclusion that NFE2L3 may involved in promoting tumorigenesis.

While prognostic clinical markers are useful, existing molecular makers still of limited value in cancers. Recent studies suggested that NFE2L3 has an ideal diagnostic and prognostic value in bladder, colorectal and renal cancers ^[8, 10, 16]^. Consistent with our findings, NFE2L3 functioned as an oncogene in PAAD and LIHC based on the fact that patients in the high-expression group had a worse prognosis than those in the low-expression group. Our results also confirmed that NFE2L3 showed ideal diagnostic and prognostic value for DSC. Therefore, NFE2L3 is expected to emerge as a novel indicator for the diagnosis and prognosis of DSC.

It is well established that cancer is a complex and heterogeneous genomic disease, and gene functions cannot be guaranteed based on the basis of standard definitions. One explanation for the aberrant function of genes in cancer is that different cellular contexts result in a wide range of pathological outcomes ^[17]^. A detailed mechanistic explanation for this problem has not yet been fully elucidated, it is worth investigating in the future.

ICIs as a mainstay of cancer treatment, have provided novel strategies and improved clinical outcomes in multiple cancer types. However, response rates to immunotherapy various owing to several critical factors, including mutation burden, tumor-infiltrating lymphocytes and regulatory checkpoint receptors^[18, 19]^ Researchers found that NFE2L3 played an important role in various immune-related pathways and immune cells. In Nfe2l3-/- mice, the levels of FOXP3 and the immune checkpoint markers cytotoxic T-lymphocyte associated protein 4 (CTLA4), TIM3, and lymphocyte activating 3 (LAG3) were increased ^[20]^. The enrichment analyses revealed that NFE2L3 was associated with various of immune-relevant pathways in kidney cancer and was related to the infiltration ratios of 17 types of immune cells in patients with kidney cancer ^[21]^. Nevertheless, the molecular mechanisms behind it remain mysterious.

The results of our study indicated a strong correlation between NFE2L3 expression and MIS in DSC. MSI is a certain phenomenon that often arises in tumors that have an underlying defect in DNA mismatch repair ^[22]^. Patients with high MSI rates indicating poor median PFS or OS ^[23–25]^. Therefore, it can be inferred that MSI could play a crucial role in shaping the progression of DSC. Relevant studies have indicated that cancers harbor high TMB means the potential to generate many neoantigens, making tumors more immunogenic for ICIs ^[26, 27]^. Expect MSI, the TMB also demonstrated the significance of the DSC. A higher TMB is normally related to an adverse outcome in advanced tumors.

According to our results, we revealed that NFE2L3 was closely correlated with immunity. It is expected to become novel immunotherapeutic target for DSC in the future. Further studies should continue to investigate the mechanisms of NFE2L3 in immunotherapy.

Our findings indicated a significant positive correlation between NFE2L3 and immune or molecular subtypes. Cellular and immune subtypes are the main factors responsible for tumor heterogeneity. Treatment elicits varied responses from tumors, owing to the genetic diversity within the tumor. The combination of immune and molecular subtypes for precisely targeted tumor therapy will also greatly enhance the current status of immunotherapeutic efficacy. Therefore, we believe that NFE2L3 plays a vital role in DSC immunotherapy.Therapeutic cancer vaccines remain a valid immunotherapy option and hold significant implications for the clinical advancement of cancer patients ^[28, 29]^. Over the past few years, there has been a significant amount of research conducted on vaccines and therapeutics specifically designed for the innate and adaptive immune systems in the field of cancer treatment ^[30, 31]^. Thus, the important functions of immune cells in tumor-specific immunity against immune cells can be understood. Our findings showed that compared with non-responders, patients who generated antibody responses to therapy exhibited prolonged DFS. We have reason to believe that NFE2L3 to be a potential target for immunotherapy in the future.

A multitude of studies have indicated that patients with immune cell infiltration tumors exhibit a more favorable response to ICI than patients with non-immunological tumors. Recent research has revealed that NFE2L3 plays a pivotal role in cancer development ^[8]^ ^[14, 32, 33]^, but the precise mechanisms have yet to be elucidated. The findings of this research indicated that DSCs experienced immune cell infiltration. The immune cell infiltration encompasses a wide range of cell types, such as Tregs, T cells, dendritic cells, NK cells, T follicular helper cells, macrophages, mast cells, and monocytes. In STAD, we observed a strong correlation between NFE2L3 expression and immune cells abundance. The research revealing the correlation of immune cell infiltration with NFE2L3 may open unique avenues for precise cancer therapeutics against cancers.

Co-expression analysis can be employed as a means to identify disease associated genes and gene functions in tumors. Chemokines combine with the cell surface chemokine receptors to perform biological functions, such as the chemotaxis, leukocyte migration, and inflammatory activities ^[34, 35]^. The causative role of chemokines is variable, and they play an important role in many diseases, such as cancer, viral infections, inflammatory, and autoimmune diseases. Researchers have found that chemokines affect anti-tumorigenic activity by regulating tumor angiogenesis and infiltration of immune cells ^[36]^. The results of our study showed a strong correlation between NFE2L3 and immune related genes or chemokine receptors. In COAD, LIHC, and STAD, there was a close correlation between NFE2L3 expression and the chemokine receptors, such as C-X3-C motif chemokine receptor 1 (CX3CR1), plexin B2 (PLXNB2), atypical chemokine receptor 1 (ACKR1), and formyl peptide receptor 2 (FPR2). Anja Schmall found that the tumor-associated crosstalk between macrophages and cancer cells via the CCR2 and CX3CR1 signaling pathways directed the lung cancer growth and metastasis ^[37]^. Circ_0013958 plays an oncogenic role in ovarian cancer by regulating the miR-637/PLXNB2 axis ^[38]^. Immune related genes, such as PD-1, PD-L1, and CTLA4, have been successfully used in tumor immunotherapy and have been shown to have a significant effect on B cell lymphomas, HCC, and non-small cell lung cancer ^[39–41]^. This study revealed novel immune related genes that are co-expressed with NFE2L3 in DSC, including TIGIT, IDO1, ADORA2A, and CD70, which can be used as immune therapy, checkpoints and biomarkers ^[42, 43]^. Therefore, combining immune-related genes or chemokine receptors with NFE2L3 may enhance the efficacy of DSC diagnosis and immunotherapy.

We conducted GO enrichment and KEGG analysis on the NFE2L3 binding proteins, we found that the Hedgehog signaling pathway is the primary target of protein catabolic processes and cell differentiation. Hedgehog signaling pathway activated tumors by driving EMT, and the inhibitor of Hedgehog signaling pathway exhibited remarkable clinical outcomes in various types of cancer^[44]^. Nevertheless, the investigation into the differential regulation of target genes by NFE2L3 remains to be investigated.

NFE2L3 is involved in the regulation of various biological and cellular processes such as cell cycle, cell differentiation or inflammatory processes ^[31]^. We performed GSEA and found that NFE2L3 was significantly associated with epigenetics, methylation modification, cell cycle regulation, calcium signaling pathway and steroid hormone biosynthesis. Consequently, the distinct manifestation of NFE2L3 assumes a crucial function in DSC through various biological functions and signaling pathways, ultimately impacting the process of tumorigenesis. Nevertheless, a comprehensive understanding of the findings would require further experimental validation.

To validate the bioinformatics findings, in vivo experiments conducted on gastric cancer cells demonstrated the role of NFE2L3 as an oncogene, facilitating the proliferation and movement of gastric cancer cells. NFE2L3 enhance tumor cell migration ability by affecting the EMT through Wnt/ β-catenin signaling pathway ^[14]^, this findings and our hypothesis are validated experimental results. More in vitro research is needed to gain a better understanding of how things work in the future.

The outcomes of these bioinformatics analyses lay the groundwork for future comprehensive investigations into the mechanisms underlying tumorigenesis and evolution. NFE2L3 was found to be aberrantly expressed in DSC, and significantly correlated with the diagnosis and prognosis of DSC patients. In addition, in vivo results suggest that NFE2L3 may act as an oncogene roles in gastric cancer through promoting the gastric cancer cells growth and migration.

## 5. Conclusions

In conclusion, we identified a specific role of NFE2L3 in DSC. The results of our study indicated that NFE2L3 was differentially expressed in cancers and was closely related to clinical features. Therefore, it has the potential prognostic and diagnostic value for DSC. In addition, NFE2L3 was closely related to immune filtration, immune subtype, and molecular subtype and was co-expressed with numerous genes and chemokines. These findings suggest that NFE2L3 has the potential to emerge as a novel therapeutic target for DSC. Additional experimental studies are required to verify these findings.

